# Metadiffusion: inference-time meta-energy biasing of biomolecular diffusion models

**DOI:** 10.64898/2026.02.10.704873

**Authors:** Hilbert Yuen In Lam, Sebastián Pujalte Ojeda, Michaela Brezinova, Josef Hanke, Xing Er Ong, Yuguang Mu, Michele Vendruscolo

## Abstract

Biomolecular function often depends on conformational ensembles, yet modern diffusion-based structure generators are biased toward the compact conformations prevalent in structural databases, limiting their ability to explore broad conformational landscapes. This work introduces metadiffusion, where an additional meta-energy biasing layer on top of diffusion steers pretrained biomolecular diffusion models through gradient-guided denoising. Without retraining, metadiffusion generates diverse conformational ensembles whose residue-level flexibility patterns closely match molecular dynamics simulations. The method supports three complementary modes: optimisation, steering to user-specified targets, and exploration via inter-sample repulsion. This approach enables controlled exploration of collective variables, enumeration of alternative binding poses across proteins, nucleic acids and ligands, and conformational ensemble generation consistent with SAXS and NMR chemical shifts. Metadiffusion thus provides a practical route to connect diffusion-based structure generation with ensemble-level, experimentally-restrained structural analysis.

## Introduction

For more than half a century, protein science has been shaped by the sequence-structure-function paradigm, in which an amino acid sequence encodes a well-defined native structure that in turn determines biological activity^1–4^. This view has been extraordinarily productive, enabling mechanistic explanation and engineering across structural biology and drug discovery, yet its very success has also revealed its limits^5,6^. Many proteins populate multiple interconverting conformations, remain partially disordered, or switch states as cellular conditions, interaction partners and post-translational modifications change^5,6^. As a result, function often emerges from a context-dependent distribution of states rather than a single structure, motivating approaches that can generate and refine structural ensembles so that conformational landscapes become a quantitative object of prediction and mechanistic interpretation^7^.

Molecular dynamics (MD) simulations have enabled extensive conformational sampling of biomolecular free energy landscapes, in particular in combination with experimental guidance^8,9^. MD sampling exhaustiveness can be further amplified through the use of enhanced sampling methods such as metadynamics where auxiliary potentials are introduced during simulations to push states into rarer, albeit physically plausible conformations that are harder to obtain from short MD simulations^10^. In addition to generating more varied states, methods such as metainference allow direct biasing of MD simulations to experimental data such as residual dipolar couplings and chemical shifts in nuclear magnetic resonance (NMR) spectroscopy^11^.

Recent advances in machine learning and biophysics have also allowed for the rapid and highly accurate *ab initio* prediction of structures as demonstrated by AlphaFold3^3^ and Boltz-2^12^. One commonality of AlphaFold3 and Boltz-2 is that they are diffusion models - machine learning models which are trained to iteratively denoise a 3D point cloud of biomolecules from random noise. AlphaFold3 and Boltz-2, on their own, however, are not well-suited to predict a diverse ensemble of structures, as they were trained on native structures in the Protein Data Bank (PDB)^13^.

Approaches have since emerged to address this problem, including flow-matching to generate conformational ensembles^14^, manipulation of the input multiple-sequence alignments (MSAs)^15,16^, or training the diffusion models directly on MD-generated ensembles and fine-tuning on experimental data^17^. While effective in certain settings, these approaches often require task-specific adaptation, and provide limited control in driving structures towards experimental observables at inference time.

Further ideas at the intersection of diffusion models and experimental measurables have been recently explored. Particle guidance has been used to enforce diversity in small molecule conformer generation^18^, and biasing away from the root mean squared distance (RMSD) of a fixed structure using sequential Monte Carlo samplers whilst constraining secondary structures was previously explored^19^. Score guidance has also been used to steer AlphaFold3 ensembles towards NMR nuclear Overhauser effects (NOEs) and cryo-electron microscopy (cryo-EM)-resolved electron densities^20^. Additionally, recent work inspired by metadynamics has introduced collective variable (CV)-based guidance into molecular diffusion models^21^. For disordered proteins specifically, models such as idpSAM incorporate biasing potentials towards radius-of-gyration (*R*_*g*_)^22^, while ExEnDiff^23^ extends diffusion-based ensemble generation on IDPs using the Str2str^24^ model to multiple experimental modalities including NMR, and cryo-EM.

Metadiffusion unifies these directions by defining a meta-energy that encodes collective-variable objectives and applying its gradients as additional updates during diffusion denoising. Conceptually, this mirrors the role of the meta-energy in metainference^11^, as it combines prior information with experimental discrepancy terms, although here it is implemented as inference-time guidance on a pretrained diffusion model rather than Bayesian sampling under an explicit force-field posterior. Metadiffusion operates directly on pretrained biomolecular diffusion models without retraining or fine-tuning, bringing together ideas from enhanced sampling, experimental ensemble refinement and score-guided diffusion within a single, composable inference-time framework. This positions metadiffusion as a general, model-agnostic approach for generating diverse, experiment-consistent conformational ensembles.

## Results

### Metadiffusion applied to Boltz-2

To illustrate metadiffusion, it is applied to the diffusion module in Boltz-2. At each denoising step, gradient-based guidance biases the intermediate atomic point cloud toward objectives defined by simple potentials, similar to previous diffusion guidance strategies^25^. Using this mechanism, metadiffusion enables: (1) matching user-specified structural descriptors such as radius of gyration (*R*_*g*_), hinge angles and solvent-accessible surface area (SASA); (2) generating highly diverse conformational ensembles and flexibility patterns that closely track MD fluctuations; (3) traversing alternative binding poses in protein-ligand and protein-nucleic-acid complexes; and (4) producing conformational ensembles consistent with experimental small-angle X-ray scattering (SAXS) data and NMR chemical shifts (**Fig. 1**).

**Figure 1.**
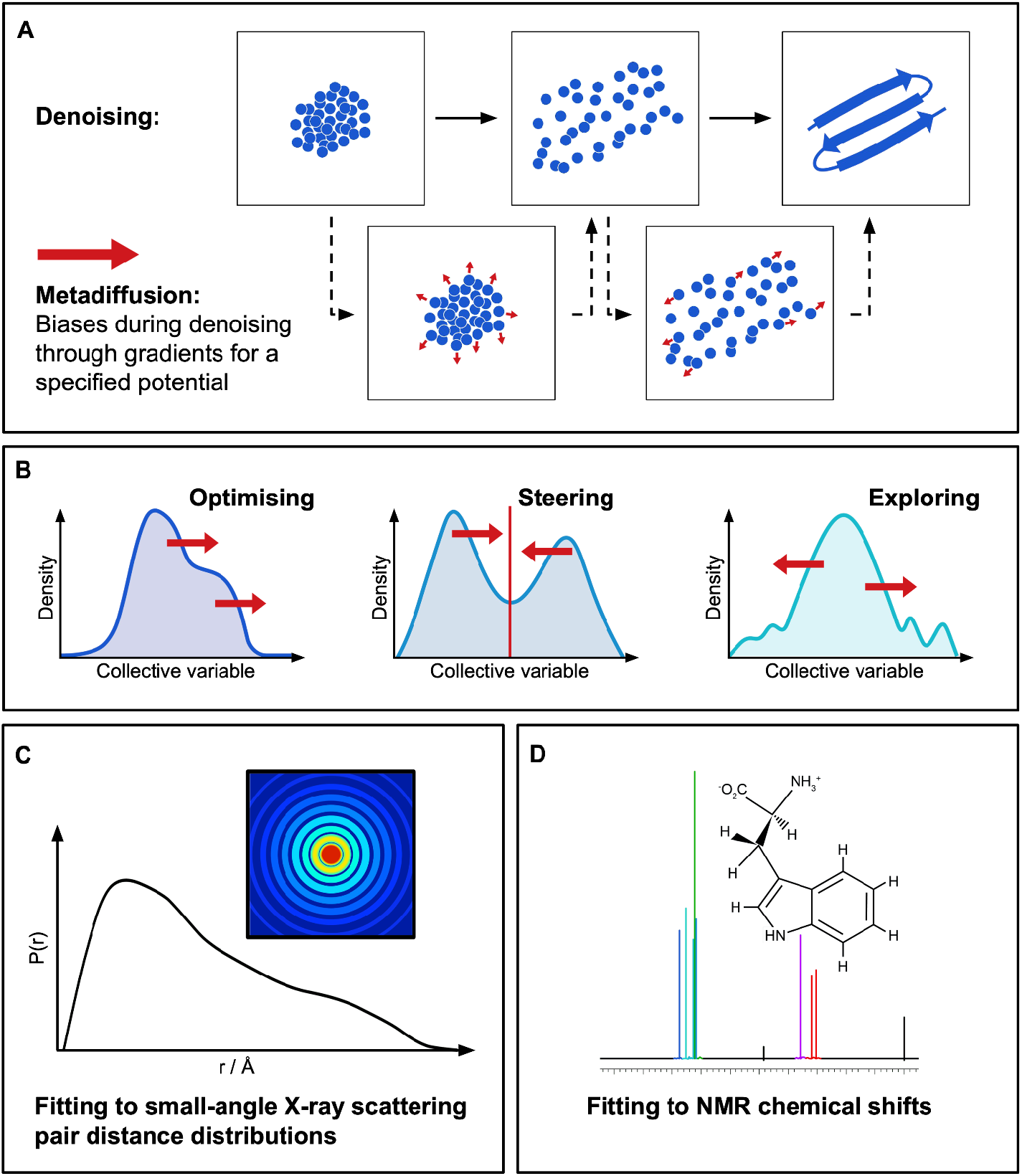
Inference-time meta-energy biasing in metadiffusion. (a) Metadiffusion applies a meta-energy guidance term during each denoising step, adding a gradient-based correction to the predicted displacement so that the atomic point cloud is steered toward a user-defined objective while remaining anchored to the learned structural prior of the model. (b) Given a differentiable CV, three primary types of biases can be injected: optimisation, in which a CV is minimised or maximised; steering - in which the biases push structures toward a user-defined value; exploring - in which the biases induce diversity in the CV. (c) Metadiffusion can be used to fit structures to SAXS pair distance distribution data. (d) The biasing can also be applied to fit NMR chemical shifts.

### Metadiffusion generates biomolecular structures controllable with user-defined structural descriptors

Minimisation and maximisation of SASA can be achieved with metadiffusion through optimisation, with side chains adopting more open and solvent-accessible positions when SASA is maximised, and pushing inward when minimised. This procedure is illustrated in the case of myoglobin (**Fig. 2a**). The steering of the *R*_*g*_ was also achieved through the use of metadiffusion - the distance between the N- and C-terminus regions increased with specification of *R*_*g*_, although at conservative metadiffusion strengths the *R*_*g*_ capped at approximately 45 Å (**Fig. 2b**). Higher strengths were able to force the *R*_*g*_ into further extrema, but at the cost of more unphysical structures (**Supplementary Table 1**). Through the introduction of Gaussian repulsion potentials between the diffusion samples (the exploration modality in metadiffusion), the monomeric MfnG of *S. drozdowiczii*, with the hinge which typically attaches to the other chain^26^ can be effectively sampled across a wide range of angles. This range is larger than that of default Boltz-2 (**Fig. 2c**). The hinge angles were calculated from the centroid of all atoms in the differently coloured regions.

**Figure 2.**
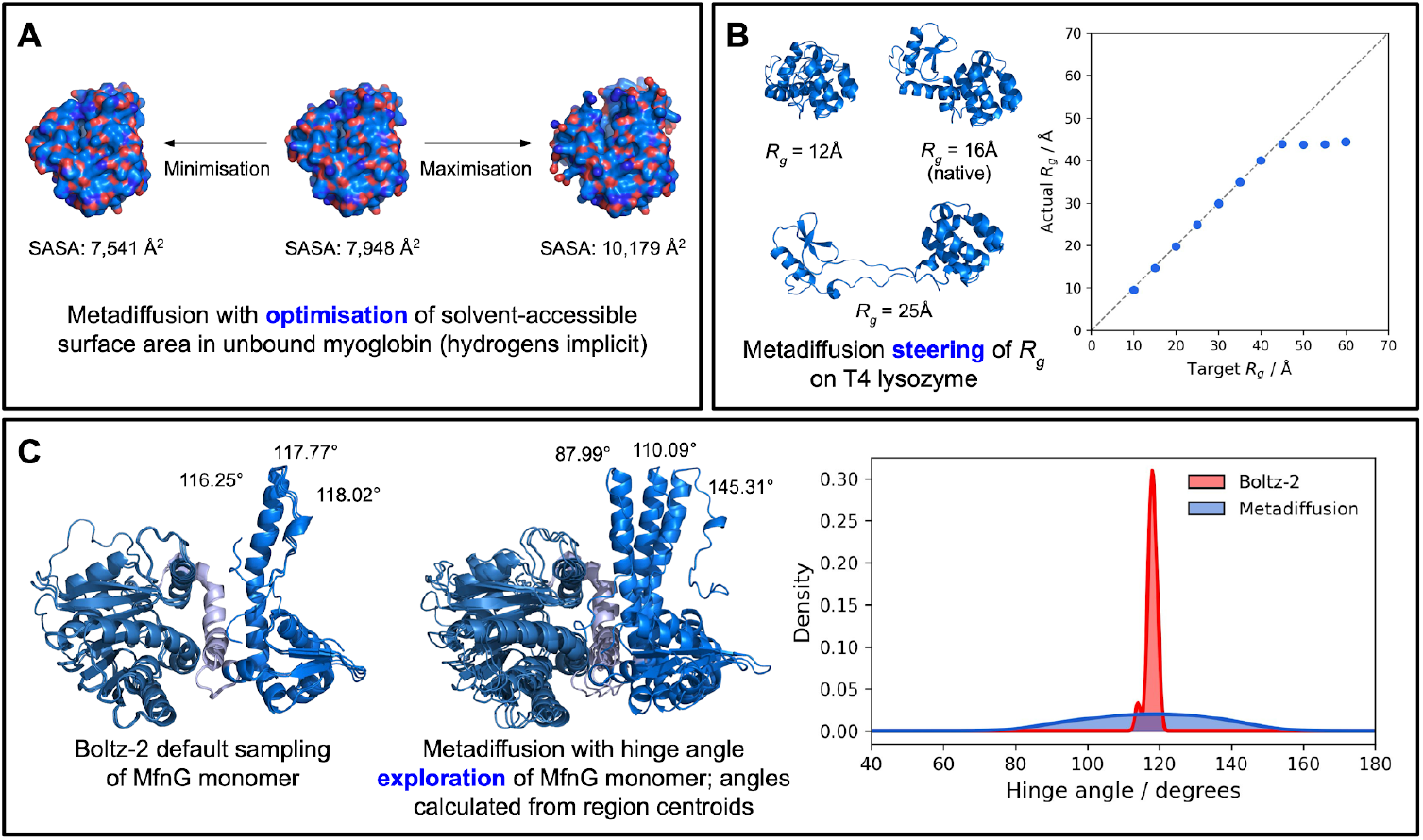
Steering collective variables with metadiffusion. (a) Optimisation, through the minimisation and maximisation of SASA in *P. macrocephalus* myoglobin. (b) Steering of T4 lysozyme *R*_*g*_. (c) Exploration of hinge angles in monomeric *S. drozdowiczii* MfnG with steering regions and vertices defined with the different colour regions, and the centroid of each differently coloured region used as the sides and vertex respectively for steering and calculation.

Through the use of the optimisation, steering and exploration modalities, a variety of other CVs can also be manipulated, including: distances, dihedral angles and shape gyration. It is also notable that high biases and/or disallowing Boltz-2 sufficient iterations to relax the structure unbiased can result in an increased number of bond breakages and steric clashes. Therefore, tuning of the bias strengths is required to achieve both metadiffusion objectives whilst maintaining physically-plausible structures. However, the majority of clashes and bond breaks can be resolved with energy minimisation (Supplementary Table 1), indicating that the method is viable to generate conformers. Energy minimisation, however, also resulted in shifts in the structure’s *R*_*g*_ values, with the lysozyme T4^27^ (which has a native *R*_*g*_ of approximately 16 Å) conformers generated at an overly compact *R*_*g*_ of 9.55 ± 0.03 Å expanding to 14.1 ± 0.1 Å (n = 8) and conformers generated at an overly extended *R*_*g*_ of 43.9 ± 1.5 Å contracting to 42.6 ± 1.5 Å (n = 8).

### Maximising the RMSD between samples in a conformational ensemble creates highly diverse conformers

Structures of different conformers of *E. coli* adenylate kinase were explored and compared with BioEmu and unbiased Boltz-2 demonstrating that metadiffusion can generate structures that are even more diverse. However, the structures generated are further away from the reference inhibitor-bound closed and open states (PDB 1AKE^28^ and 4AKE^29^, respectively). The diversity of structures also scales monotonically to the strength of bias applied in both protein and nucleic acid regimes (**Fig. 3b, c**). These results suggest that conformational diversity can be controlled. Since the biasing can be applied simultaneously to multiple groups (i.e. different chains in a structure such as a ligand), different poses of the small molecule drug risdiplam were elucidated for an RNA duplex^30^. The generated structures controllably exceed the diversity generated resulting from NMR structural determination (**Fig. 3b**).

**Figure 3.**
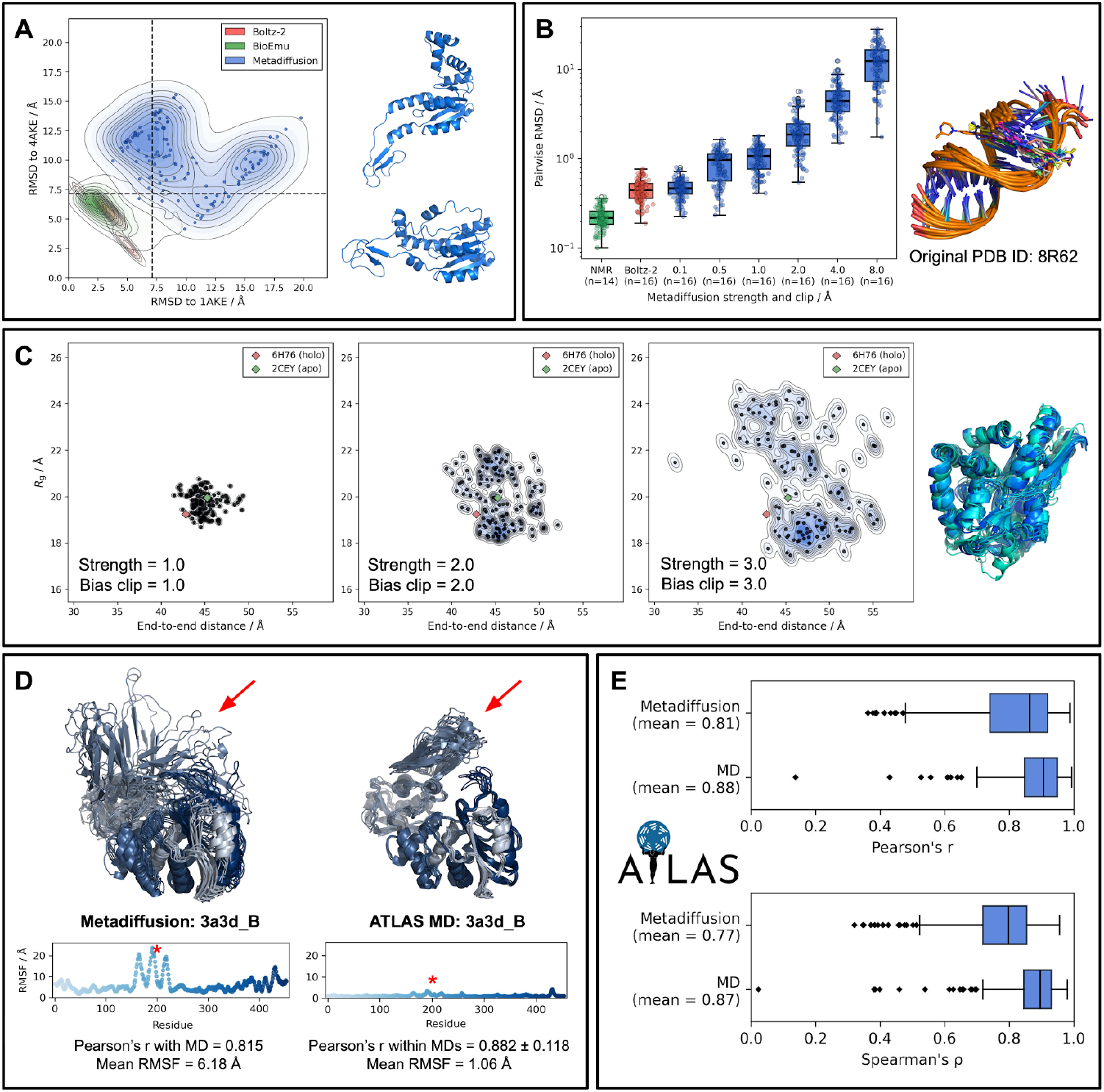
Diversity generation through RMSD-based exploration. (a) *E. coli* adenylate kinase RMSD with reference to the ligand-bound closed (1AKE^28^) and open (4AKE^29^) states on the PDB with 128 conformers generated for metadiffusion and Boltz-2, and 4,000 structures generated in BioEmu using the Heun denoiser. Individual structure RMSDs between the two states from metadiffusion are shown as dots. RMSDs are calculated using a single-step Kabsch alignment. (b) Biasing of risdiplam bound to RNA duplex using pairwise RMSDs between samples with increasing strengths and metadiffusion bias clips showing controllable diversity through metadiffusion parameters, with box-and-whisker plots indicating the pairwise RMSD between generated structures. 128 structures were generated with different strength and bias clip parameters. (c) Stronger biases and looser clipping allow the method to push the model further away from its learned prior, thereby generating more diverse structures; this is illustrated for the native state of *H. influenzae* SiaP^62^. (d) Structures of metadiffusion-generated compared with 100 ns MD simulation of single-chain *H. influenzae* dacB starting from the native structure (3A3D^63^). Plots show the RMSF in each C_α_ versus MD, with emphasis on the highly flexible domain. (e) Box-and-whisker plots of the Pearson’s R and Spearman’s ρ of RMSF correlations with 256 randomly-selected MD simulations of 100 ns each from the ATLAS dataset^31^.

### Biased ensembles fluctuate similarly to that in MD

Diverse structures generated by maximising the pairwise RMSD between samples in metadiffusion correlate strongly with 100 ns MD simulations in the ATLAS dataset^31^. This is indicated by the mean Pearson’s R of 0.81 when comparing the RMSF of the C_α_ of 256 randomly-selected structures covered by ATLAS. Since ATLAS ran the MD simulations in triplicates, it is noteworthy that between the MD simulations themselves, an average Pearson’s R of 0.88 was achieved, indicating that fluctuations from metadiffusion are close to the reproducibility ceiling. However, the mean RMSF per protein predicted by metadiffusion only correlates with a Pearson’s R of 0.65 with the mean RMSF per protein in MD simulations, indicating the magnitude of fluctuation between the proteins differs. The exact Pearson’s R correlation metrics and protein identities selected from the ATLAS dataset are available in **Supplementary File 1**.

### Diverse structures are reasonable and can be effectively improved with energy minimisation

Structures generated by maximising the pairwise RMSD in ATLAS have occasional bond breaks and steric clashes, but can be effectively resolved with energy minimisation with an implicit solvent force field (**Supplementary Fig. 1**). Following minimisation, very few of the 256 × 16 structures displayed peptide bond lengths of > 1.7 Å and steric clashes were significantly reduced by 89.9%.

### Different docking poses of proteins, nucleic acids, and ligands can also be obtained by biasing the ligand whilst aligning to the structure

In doing the Kabsch alignment during the denoising process, the RMSD of the ligand can be calculated relative to one group (such as a chain) while aligned to another. By aligning one chain and applying biases solely to another, the relative binding poses between the two can be explored. This is effectively holding one component fixed while sampling conformational diversity in the other. For example, different poses of the H-Ras and Raf kinase RBD complex can be resolved^32^ (**Fig. 4a**) and DNA ligases can be mapped to different binding conformations with DNA^33^ (**Fig. 4b**). Similarly, a ligand can be biased which effectively allows Boltz-2 to co-fold the protein with a single ligand in different conformations in the same pocket (**Fig. 4c**) and also in different pockets (**Fig. 4d)**, as demonstrated by furosemide in human serum albumin. Interestingly, even though only the ligand receives the biases during this process, the protein also adopts slightly different conformations, especially locally, to accommodate the ligand. This potentially provides an advantage over traditional docking, as the protein flexibility as a whole is considered, and not just local flexibility as per flexible docking.

**Figure 4.**
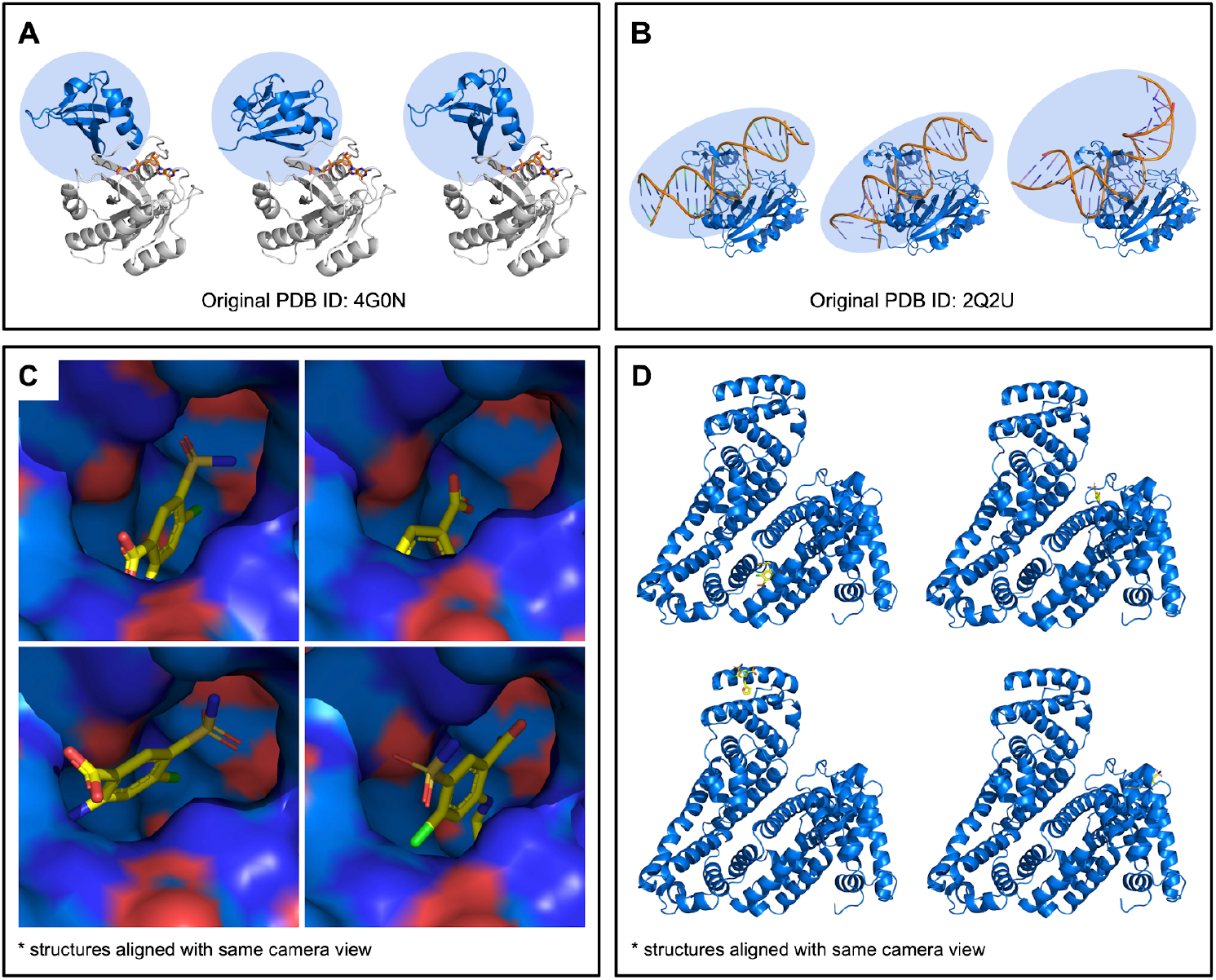
Exploring alternative binding poses with metadiffusion. (a) Biasing of the *H. sapiens* RBD of Raf kinase whilst aligned to H-Ras enumerates different binding poses of the RBD of Raf kinase. (b) Chlorella virus DNA ligase bound to DNA product in different conformations, generated by aligning to the protein whilst applying diversity biases only to the nucleic acid chains. (c) Connolly surface of furosemide in a local pocket on human serum albumin generated in one pass of metadiffusion with high strength by aligning to the protein but only biasing the ligand. The protein is aligned and superimposed in the images, and shown with the same camera view. (d) Multiple different docking sites and poses of furosemide on human serum albumin as generated by metadiffusion.

### Ensembles can be biased with metadiffusion to fit experimental observables in SAXS and NMR

Steering of class V GTP aptamer in complex with GTP to the SAXS pair distance distribution led to lower Wasserstein-1 distance between the metadiffusion-generated ensembles than the unbiased Boltz-2 structures to the experimental SAXS P(r) curve^34^ (**Fig. 5a**). With steering from NMR chemical shifts in *H. sapiens* calmodulin^35^, Pearson’s R in ^13^C_α_ increased from 0.66 to 0.97 and ^13^C_β_ increased from 0.97 to 0.99 for n = 16 structures with and without SAXS steering respectively. Similarly, the structures can be steered towards SAXS^36^ concurrently with the chemical shifts, reducing the final Wasserstein-1 distance as compared to unbiased Boltz-2 generations (**Fig. 5b**). It is also notable that the pair-RMSD dissimilarity was maximised in the generation of these structures, leading to highly-diverse structures which, as a whole, fit to the ensemble NMR and SAXS (**Fig. 5a, b**).

**Figure 5.**
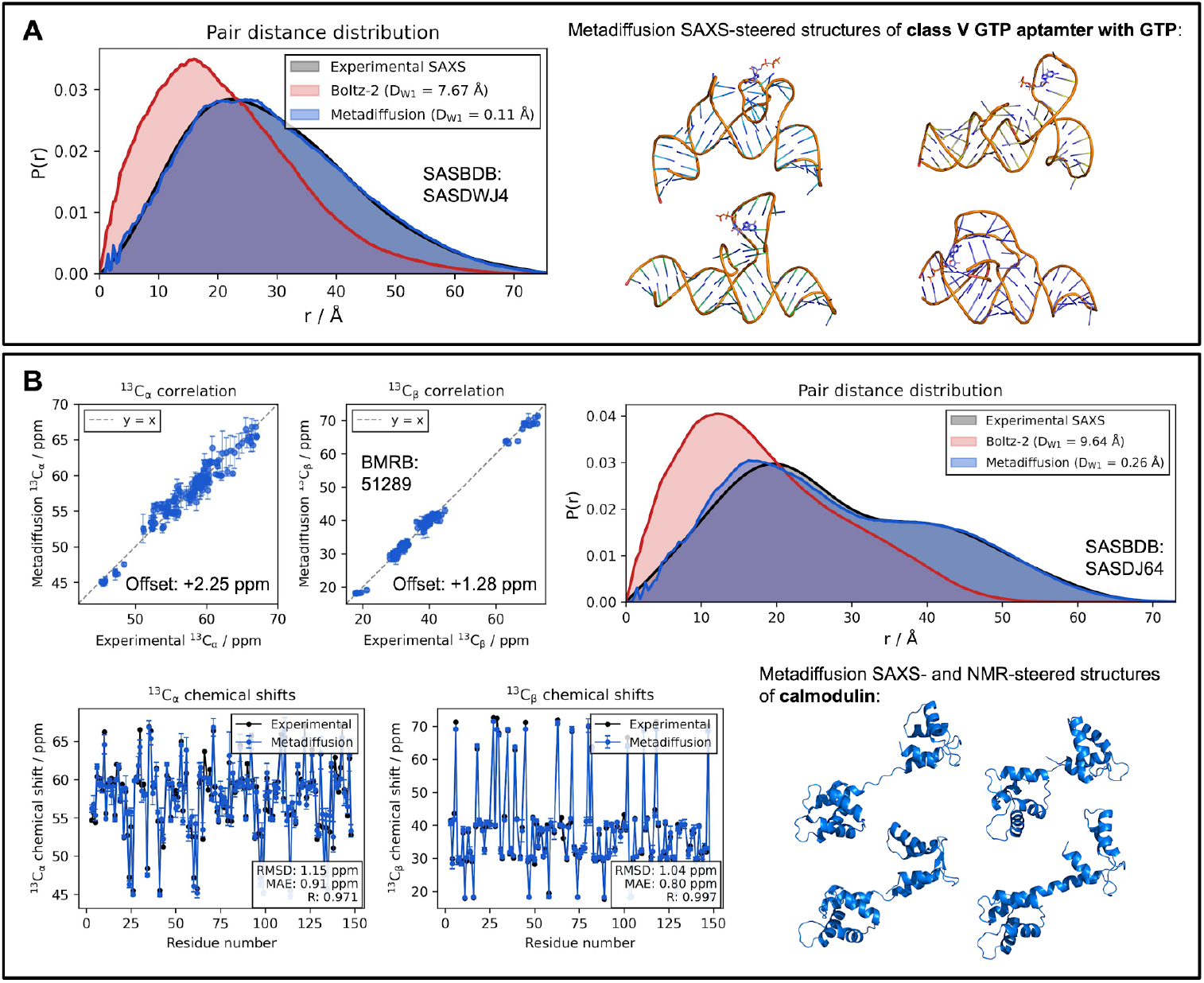
Steering conformational ensembles to experimental SAXS and NMR data. (a) Experimental SAXS pair distance distribution of class V GTP aptamer in complex with GTP molecule (SASBDB: SASDWJ4^34^) with ensemble-averaged pair distance distribution of unbiased Boltz-2 generation (n = 8) and SAXS-fitted structures with metadiffusion (n = 8). (b) Chemical shifts of *H. sapiens* calmodulin (BMRB: 51289^35^) and ensemble-averaged CheShift-2 calculated chemical shifts of SAXS and NMR-fitted metadiffusion structures. Experimental SAXS pair distance distribution (SASBDB: SASDJ64^36^) and ensemble-averaged pair distance distribution of unbiased Boltz-2 structures (n = 16) and fitted structures with metadiffusion (n = 16) are also shown.

### Speed of metadiffusion

Metadiffusion speed varies heavily by the CV used and the number of biases added. For maximising the pairwise RMSD between samples, the computational complexity is O(N^2^), with N being the number of samples, as reflected in **Supplementary Fig. 2**, where the 76-residue ubiquitin was used^37^. CVs such as *R*_*g*_ or distances take significantly less time with O(N) compute, whereas SAXS- and NMR-steering take longer due to them necessitating the pairwise distances of all atoms be calculated at each biasing step and the CheShift-2 algorithm for chemical shift calculation being run respectively.

## Discussion

The development of diffusion-based structure prediction models represents a major advance in structural biology, yet their focus on single, compact conformations inherited from crystallographic training data leaves conformational dynamics largely unaddressed. Metadiffusion bridges this gap by introducing inference-time meta-energy guidance that steers pretrained diffusion models toward diverse, physically plausible ensembles without retraining. By leveraging the rich structural priors these models have learned from experimental data and combining them with gradient-based biasing through collective variables, metadiffusion creates a practical synthesis of machine learning efficiency and physics-driven interpretability. This approach positions conformational ensemble generation not as a separate modeling task requiring specialised training, but as a controllable property of inference that can be adapted to experimental constraints and mechanistic questions on demand^7^.

Because metadiffusion applies inference-time meta-energy guidance to pretrained biomolecular diffusion models, it is model-agnostic in principle. Metadiffusion can be applied to other generative diffusion-based machine learning protein folding models such as AlphaFold3^3^, OpenFold^38,39^, Chai-1^40^, or IDPFold2^41^, without requiring any retraining or fine-tuning of these models.

Conceptually, metadiffusion parallels metainference, a method in which ensemble-consistent experimental biases are introduced to MD simulations through Bayesian modelling^11^. In metadiffusion, however, the MD engine which integrates Newton’s equations of motion is swapped out with a diffusion-based generative machine learning model, allowing ensembles to be generated with a learned structural prior rather than an explicit force field.

A key advantage of metadiffusion is its ability to generate rare, metastable states at low computational cost, orders of magnitude faster than microsecond-scale MD simulations. By guiding the diffusion process and maximising the pairwise dissimilarity between structures according to a controllable meta-energy function, metadiffusion efficiently constructs physically-plausible conformational hypotheses. In this way, metadiffusion can be used to steer biomolecules to differentiable structural properties. The steering of conformational ensembles to SAXS and NMR data also allows for better reconciliation between experimental observables and computationally-generated structures.

In this view, by biasing degrees of freedom in a molecular engine, the generated ensembles are consistent with the statistical physics implicitly encoded by the underlying model. This is inherently a double-edged sword of diffusion-based generative models used in this work. The base model captures the biomolecular distributions in a way that might be hard to model with force fields and traditional methods, but similarly may be unsuitable in cases which are less represented in its training data. This is likely more pronounced in rarely-seen proteins - which previous work has shown that the models do not predict as well because they are out-of-distribution^42,43^. This limitation, however, can be mitigated with even more rigorous models trained on more data.

Several practical limitations of metadiffusion warrant consideration. First, the method requires tuning of guidance weights and may be hyperparameter sensitive, at least in the implementation reported in this work. Secondly, the structures generated are fundamentally limited by the expressivity of the base model, as diffusion biases can be argued as compromising the fidelity of the learned prior^44,45^. Thirdly, whilst metadiffusion is effective in generating diverse structures, it does not inherently provide thermodynamic weighting. Methods that use MD to further explore and thermodynamically weight the conformational space starting from generated structures as seeds may be helpful^46^ to obtain free energy landscapes and potentially ligand binding paths^47,48^. Lastly, for heterogeneous systems such as disordered proteins, since the conformational landscapes are vast and training data sparse, the utility of metadiffusion may be constrained until base models are better able to represent these systems.

Overall, metadiffusion occupies an intermediate space between a physics-driven and data-driven generative modelling. This framework combines the efficiency and flexibility of learned models with physically-interpretable CVs and guidance mechanisms.

Future work will aim to further integrate physics-driven and data-driven approaches, uniting the bottom-up paradigm of force fields - where interactions are explicitly defined and parameterized - with the top-down paradigm of machine learning, where structural priors are learned from experimental data. In the meantime, metadiffusion provides a fast, practical, and extensible approach that can explore conformational landscapes and bridge experimental observables with structure-generation models, paving the way for a better-informed understanding of biomolecular conformational landscapes.

## Methods

### Implementation

Metadiffusion is implemented entirely through the addition of biases on top of Boltz-2 during denoising. As such, it relies entirely on Boltz-2’s trunk and diffusion model with no additional training or fine-tuning of the base model. Implementation of all CVs and the calculation of their corresponding gradients is performed in PyTorch 2.9.1+cu128.

### Introducing biases in diffusion

Biases are introduced during the denoising process of the Boltz-2’s diffusion model into the atomic point cloud directly. Typically, the biases are applied for the first 70-90% of the denoising process, and the structure is allowed to denoise unbiased for the remaining steps to enable it to settle into physically valid states. Biases are applied to all atoms at specifiable intervals. The displacement vectors from Boltz-2’s denoising process and biases are summed before being applied simultaneously in each iteration. Biases are calculated using derivatives from CVs, and backpropagated into the atomic point cloud. During the denoising process, at each biasing iteration, metadiffusion biases are calculated twice - once before and once after the denoising step. These biases are then combined with Boltz-2’s displacement vector and applied simultaneously. The exact biases and configurations used can be found in the code examples.

### Optimisation

Optimisation pushes a CV towards higher or lower values without a specific target. The energy function is

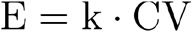

where k is the strength, and CV is the user-specified collective variable (i.e. *R*_*g*_).

### Steering

Steering biases the structure towards a user-defined value for a collective variable. It is calculated using a harmonic loss

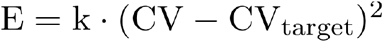

where E is the final energy, k is the strength, and CV is the user-specified collective variable.

### Exploration

Exploration attempts to create ensembles that explore a CV. A Gaussian RBF is used to push the samples in an ensemble away from each other

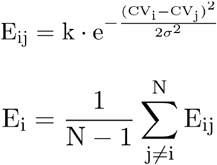

where E is the final energy, k is the strength, CV_i_ or CV_j_ is one sample’s CV value, σ is a user-defined spread of the Gaussian RBF, and N is the number of pairs. The formula ensures that the repulsion is multiplicity-invariant and a different number of samples in a diffusion run will produce the same amount of force.

### CV normalisation and noise clipping

In order to provide for meaningful steering, the gradients from all CVs presented in this work are L2 normalised before any optimisation, steering or exploration. This is to ensure that different CVs can generally be scaled with similar strength values.

### Energy minimization

Energy minimisation for generated structures was performed with OpenMM 8.4^49,50^ using the ff14SB force field^51^, implicit Generalised Born solvent (GB-Neck2)^52^, and constrained H-bonds. Hydrogen atoms were also added using PDBfixer^49,50^ to metadiffusion-generated structures at pH 7.0 to allow proper minimisation.

### Peptide bond breakage and steric clash determination

Peptide bond breaks are defined by the C-N bond exceeding 1.7 Å. For steric clash detection, all heavy atom pairs were considered, excluding those within the same or adjacent residues of the same chain. A clash was counted when the interatomic distance fell below the sum of the two van der Waals radii minus a 0.4 Å tolerance.

### RMSD pair

In the RMSD pair-biasing, all RMSDs between the *i* and *j* structures are computed with single-iteration Kabsch alignment. The upper triangle RMSDs are then maximised as per the optimisation algorithm.

### BioEmu conformer generation

BioEmu conformers were generated using the Heun denoiser with 100 steps.

### Other CVs

The SASA CV is calculated using the LCPO algorithm^53^, with the SASA reported in the results calculated using the BioPython 1.84 PDB.SASA module for orthogonality. The angle CV is calculated from the centroid of all diffused atoms from a region, and *R*_*g*_ is calculated from the positions of the atoms during denoising and ignoring hydrogen atoms (as they are not included in Boltz-2 denoising).

### SAXS fitting

SAXS fitting was performed by minimising the Cramér distance^54^ from an input pair distance distribution (P(r)), computed across all atoms in the structure. Since Boltz-2 does not generate hydrogens in its diffusion process, these are ignored in the pairwise calculation. The distance distribution is binned into 32 bins before the Cramér distance is calculated and minimised. The re-binning is to ensure that different SAXS profiles are divided similarly, and to prevent diminished gradients from too many bins. Although SAXS measurements reflect distances between all electrons, this implementation approximates the scattering signal using interatomic distances, following a common atomistic approximation as employed by previous work^55,56^.

### SAXS pair distances used

The SAXS profile for class V GTP aptamer in complex with GTP was obtained from the Small-Angle Scattering Biological Data Bank (SASBDB)^57^ with the ID of SASDWJ4^34^ as a P(r) pair distance distribution. Similarly, the profile for calmodulin was obtained from SASBDB with the ID of SASDJ64^36^.

### NMR chemical shift fitting

NMR chemical shifts for C_α_ and C_β_ were calculated using a faithful PyTorch implementation of CheShift-2^58^. CamShift^59^, the default method in PLUMED 2^60^, was not used as contributions from hydrogens would be necessary, and Boltz-2 does not generate hydrogens in its denoising process. Fitting was performed by minimising a concordance correlation coefficient distance. An automatic offset was applied between mean experimental and CheShift-2 predicted shifts at each biasing iteration, calculated by taking the mean of the experimental shifts subtracted by the mean of the calculated shifts. The offset is calculated for both C_α_ and C_β_. This accounts for systematic offsets between CheShift-2’s DFT reference frame and experimental DSS reference without requiring manual calibration.

### NMR spectra used

The NMR spectra for *H. sapiens* calmodulin can be obtained from the Biological Magnetic Resonance Bank (BMRB)^61^ ID 51289^35^.

## Acknowledgements

The authors thank Adelie Louet and Ellis Baker for helpful conversations. Claude Code (Opus 4.5 / Sonnet 4.5) was employed in the implementation of this work.

## Code availability

Code is available at https://github.com/Chokyotager/Boltz-Metadiffusion.

## Funding

This work is supported by UKRI (10059436, 10061100 and 10138075). Hilbert Yuen In Lam and Sebastián Pujalte Ojeda are supported by the Una and Derek Finlay scholarship. Part of this work was performed using resources provided by the Cambridge Service for Data Driven Discovery (CSD3) operated by the University of Cambridge Research Computing Service (www.csd3.cam.ac.uk).

**Supplementary Figure 1.**
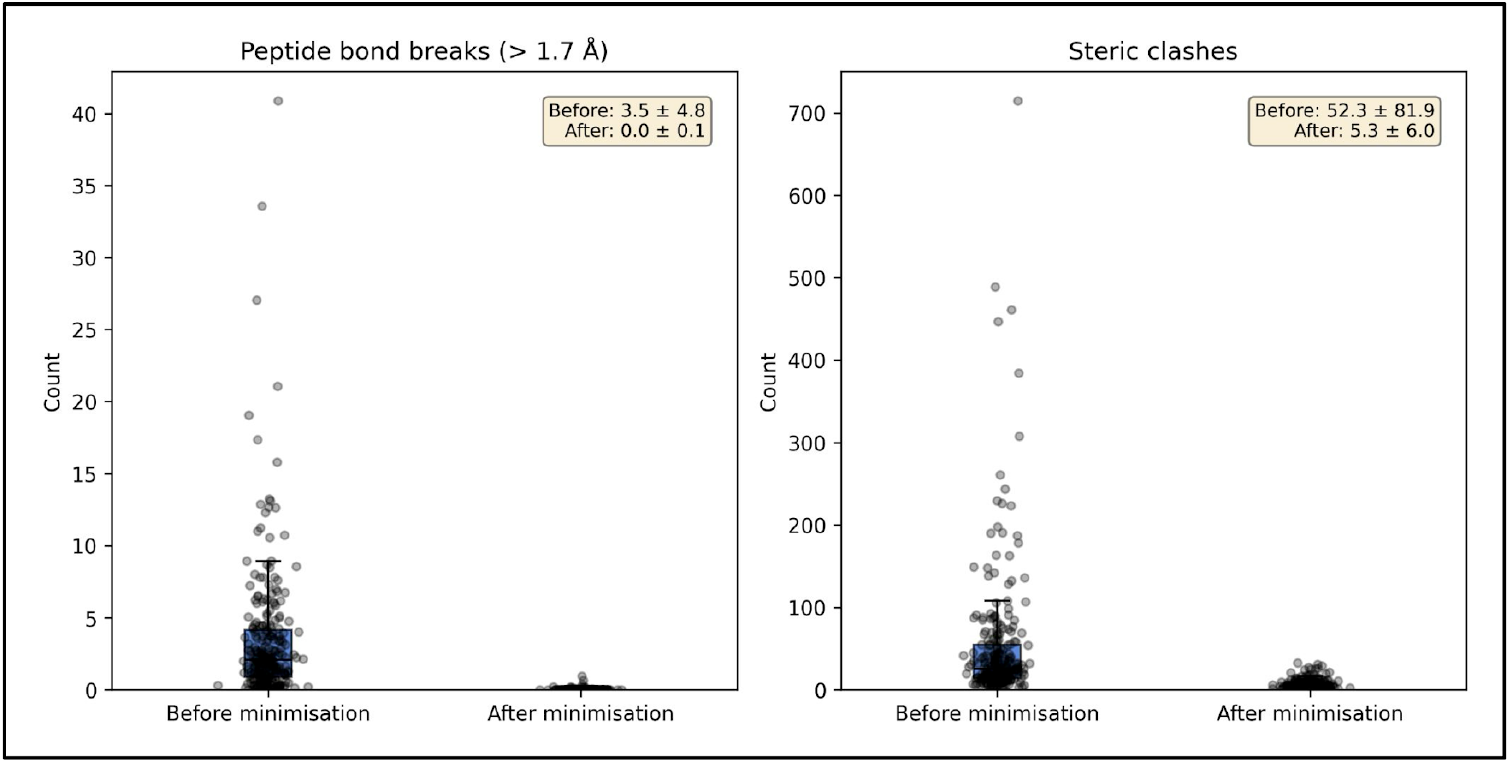
Structural quality of ATLAS-derived conformational ensembles before and after minimisation. 16 structures were generated for 256 ATLAS proteins at strength=3 and bias clip=3 to maximise pairwise RMSD between diffusion samples. Peptide bond breaks are defined as > 1.7 Å and steric clashes are determined by van der Waals radii. Each point indicates the mean peptide bond breaks/steric clashes averaged across 16 structures generated by metadiffusion in a single run. Minimisation was performed with the ff14SB force field and implicit solvent with GB-Neck2.

**Supplementary Figure 2.**
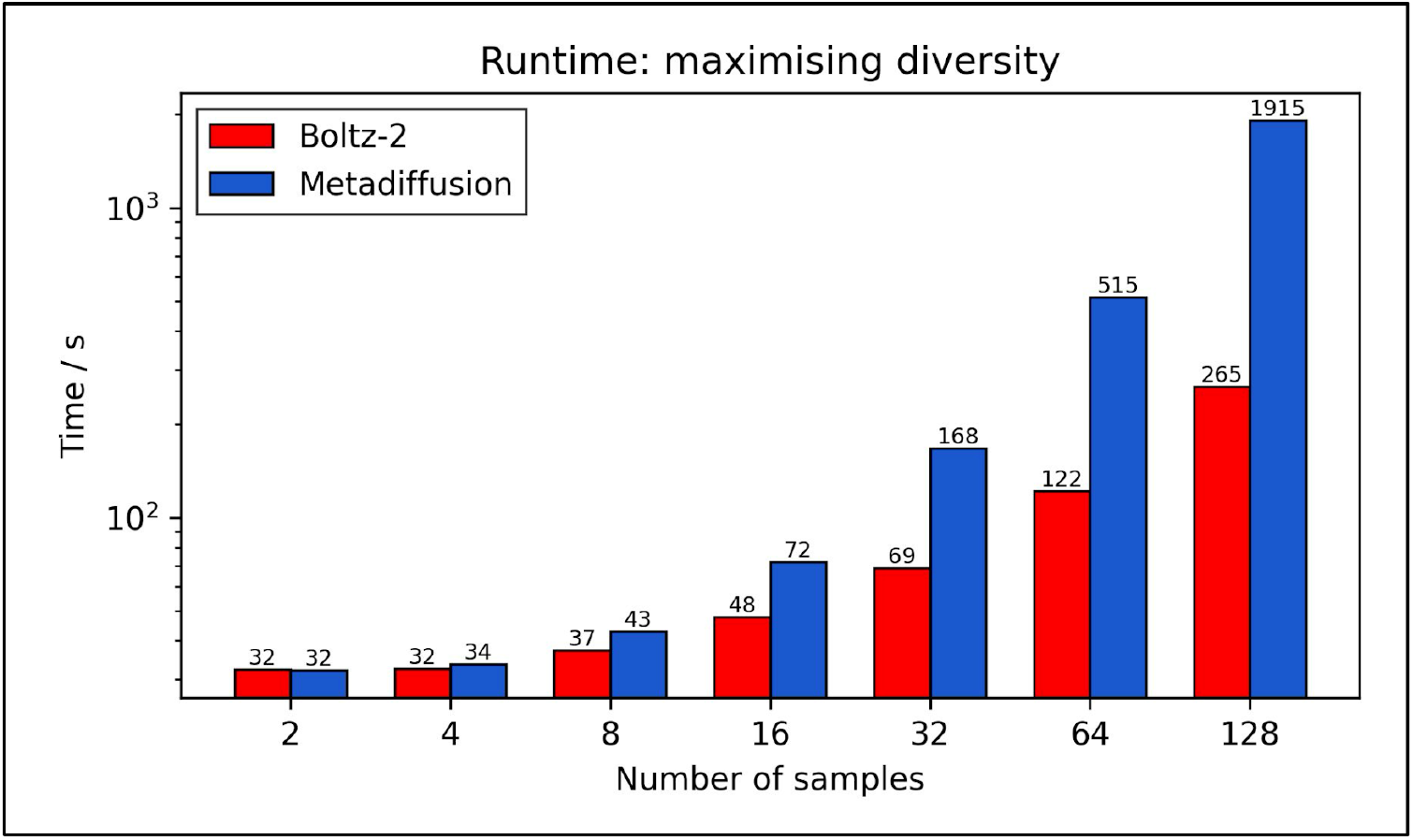
Computational cost of metadiffusion as a function of conformational ensemble size. Time taken to generate ensembles of different numbers of samples of ubiquitin (76 amino acids) generated with maximising the RMSD between samples in metadiffusion versus unbiased Boltz-2 benchmarked on an Nvidia RTX™ 3090 GPU.

**Supplementary Table 1.** Effect of energy minimisation on structural quality of metadiffusion-generated structures.

| Target $R_g$ / Å | Peptide bond breaks before minimisation | Peptide bond breaks after minimisation | Steric clashes before minimisation | Steric clashes after minimisation |
| --- | --- | --- | --- | --- |
| 10 | 0.0 ± 0.0 | 0.2 ± 0.4 | 4,602.0 ± 119.0 | 155.8 ± 281.4 |
| 12 | 0.0 ± 0.0 | 0.0 ± 0.0 | 1,654.0 ± 64.2 | 25.8 ± 25.6 |
| 14 | 0.0 ± 0.0 | 0.0 ± 0.0 | 355.8 ± 56.6 | 3.2 ± 1.3 |
| 15 | 0.0 ± 0.0 | 0.0 ± 0.0 | 74.1 ± 20.7 | 3.4 ± 4.4 |
| 16 | 0.0 ± 0.0 | 0.0 ± 0.0 | 13.2 ± 3.8 | 1.8 ± 0.8 |
| 20 | 0.0 ± 0.0 | 0.0 ± 0.0 | 1.6 ± 1.5 | 1.8 ± 1.1 |
| 25 | 0.0 ± 0.0 | 0.0 ± 0.0 | 0.5 ± 0.7 | 1.5 ± 0.7 |
| 30 | 0.0 ± 0.0 | 0.0 ± 0.0 | 0.9 ± 1.2 | 0.8 ± 0.7 |
| 35 | 0.6 ± 1.0 | 0.0 ± 0.0 | 0.6 ± 0.7 | 0.8 ± 0.7 |
| 40 | 5.2 ± 3.0 | 0.0 ± 0.0 | 0.1 ± 0.3 | 0.5 ± 0.7 |
| 45 | 21.0 ± 7.7 | 0.0 ± 0.0 | 0.0 ± 0.0 | 0.1 ± 0.3 |

**Supplementary File 1. ATLAS trajectory IDs used to calculate correlation with metadiffusion and individual Pearson R values**. RMSD biases were calculated with strength=3 and bias clip=3.

